# Overcoming Hemophilia A Gene Therapy Limitations with an Enhanced Function Factor VIII Variant

**DOI:** 10.1101/2024.05.16.594568

**Authors:** Anna R. Sternberg, Cristina Martos-Rus, Robert J. Davidson, Xueyuan Liu, Lindsey A. George

## Abstract

Durable factor VIII (FVIII) expression that normalizes hemostasis is an unrealized goal of hemophilia A adeno-associated virus (AAV)-mediated gene therapy. Trials with initial normal FVIII activity observed unexplained year-over-year declines in expression while others reported low-level, stable FVIII expression inadequate to restore normal hemostasis. Here we demonstrate that mice recapitulate FVIII expression-level-dependent loss of plasma FVIII levels due to declines in vector copy number. We show that an enhanced function FVIII variant (FVIII-R336Q/R562Q; FVIII-QQ), resistant to inactivation by protein C, normalizes hemostasis at below-normal expression levels without evidence of prothrombotic risk in hemophilia A mice. These data support that FVIII- QQ may restore normal FVIII function at low-levels of expression to permit durability using low AAV vector doses to minimize dose-dependent AAV toxicities. This work informs the mechanism of FVIII durability after AAV gene transfer and supports that incorporating the FVIII-QQ transgene may safely overcome current hemophilia A gene therapy limitations.

## Introduction

Hemophilia A (HA) is an X-linked congenital bleeding disorder due to a deficiency in (f)actor VIII cofactor function that is essential for enhancing FIX catalytic activity to amplify coagulation and prevent blood loss after vascular injury^1^. Clinical manifestations include life-threatening hemorrhage and recurrent joint bleeding with disabling arthropathy^2^, which are predicted by circulating plasma FVIII activity. Patients with severe HA (FVIII activity <1% of normal) have frequent and spontaneous hemorrhage that occurs less commonly in moderate HA patients (1-<5% of normal). The mild HA (5-<40% of normal) phenotype is heterogenous, but FVIII activity >15% protects against spontaneous joint bleeds^3^. The current standard-of-care is recurrent intravenous FVIII infusion or subcutaneous administration of a FVIIIa-mimetic antibody, emicizumab^4,5^. While effective, these therapies still require recurrent administration. Factor VIII gene therapy holds the promise of a one-time therapy to correct phenotype.

Current HA gene therapy efforts predominantly use adeno-associated virus (AAV)-mediated gene addition to exogenously express B-domain deleted FVIII in hepatocytes^6–9^. First-generation clinical trial data outline a series of limitations that include, among others, an AAV dose-dependent anti-AAV cellular immune response resulting in loss of transgene expression in at least 1 trial^6^, inadequate FVIII expression to correct phenotype, and unexplained multi-year declines in FVIII expression^6,7,10–13^. While the only licensed HA vector (valoctocogene roxaparvovec) initially achieved normal/near-normal FVIII activity, FVIII expression declined by almost half year-over-year such that median FVIII activity fell into the range of low mild to moderate HA within 5 years post-vector^7,10,12^. In contrast, stable FVIII expression has been observed in the range of moderate or low mild HA^6,9,14^, which is at or below the relative FVIII equivalency imparted by emicizumab prophylaxis^15^. These data demonstrate that AAV gene addition can impart durable FVIII expression, albeit possibly only at low levels of expression.

To overcome existing limitations, multiple novel approaches are in development for HA gene therapy^16^. One possibility is to express an enhanced function FVIII variant to normalize hemostasis at low plasma FVIII concentration to permit expression durability. In addition to overcoming HA gene therapy-specific limitations, the general advantage of an enhanced function protein variant is exemplified by the success of second-generation hemophilia B (HB) gene therapies that universally employ a gain-of-function FIX variant, FIX-Padua^17–19^. In the absence of a naturally occurring enhanced function FVIII variant, rationally engineering FVIII to bypass mechanisms of activated FVIII (FVIIIa) regulation is an attractive approach^20–22^. Factor VIIIa is inactivated by spontaneous A2-domain dissociation or by proteolytic cleavage by activated protein C (APC). We previously generated a FVIII variant with Arg to Gln mutations at the two established APC cleavage sites R336 and R562 (FVIII-R336Q/R562Q or FVIII-QQ)^20,23,24^. Factor VIII-QQ *in vitro* activity is analogous to wild-type FVIII and has normal A2-domain dissociation kinetics but is resistant to APC-mediated proteolytic inactivation. As a result, in recombinant protein studies, FVIII-QQ demonstrated 4-5-fold greater *in vivo* hemostatic function relative to FVIII-WT across multiple injury models in HA mice^20^. Here we investigate the durability, efficacy and safety of using AAV-mediated expression of FVIII-QQ as a second-generation approach to HA gene transfer to overcome current limitations.

## Results

### Durable FVIII expression is dependent on plasma antigen levels

Clinical trial data thus far suggest that sustained FVIII expression above the range of mild HA may not be feasible^6,7,10,12,14^. To investigate the mechanism of durable FVIII expression post AAV-mediated gene transfer, HA/C57BL/6 CD4-knockout (HA/CD4KO) mice were treated with a codon optimized AAV8 vector (CO1) to express B-domain deleted FVIII (FVIII-SQ^30^, herein FVIII-WT) or FVIII-QQ at variable plasma FVIII concentrations. These experiments were also designed to assess if there were any differences in longitudinal pharmacokinetics of FVIII-WT and FVIII-QQ expression. HA/CD4KO mice were used because immune-competent C57BL/6 mice mount a robust antibody response to human FVIII expression^25^. Animals were assigned to the mild HA (0.05 to <0.4 nM), normal (0.4 to <1.5 nM) and elevated FVIII (≥1.5 to 3 nM) cohorts based on their steady-state week 8 plasma FVIII antigen (**Fig. 1a,b** and **Extended Data Table 1**). Factor VIII expression pharmacokinetics were determined up to 72 weeks. Plasma FVIII antigen at week 8 versus week 72 did not significantly differ in the mild HA and normal FVIII cohorts, but was significantly reduced at week 72 for the elevated FVIII-WT and FVIII-QQ cohorts (**Fig. 1a,b**). Combining FVIII-WT and FVIII-QQ data again demonstrated a significant reduction in week 72 versus week 8 antigen among the elevated FVIII cohort only (**Fig. 1c**). Analysis of FVIII antigen values over time indicated that declines in expression began >6 months after AAV treatment (**Fig. 1d**) and did not correlate with alanine aminotransferase values (data not shown). The data demonstrated that elevated FVIII negatively impacts durability for both FVIII-WT or FVIII-QQ expression. These findings are the first to document plasma FVIII antigen-dependent expression durability in mice and are qualitatively analogous to human clinical trial data^6,7,10,13,14^.

**Figure 1:**
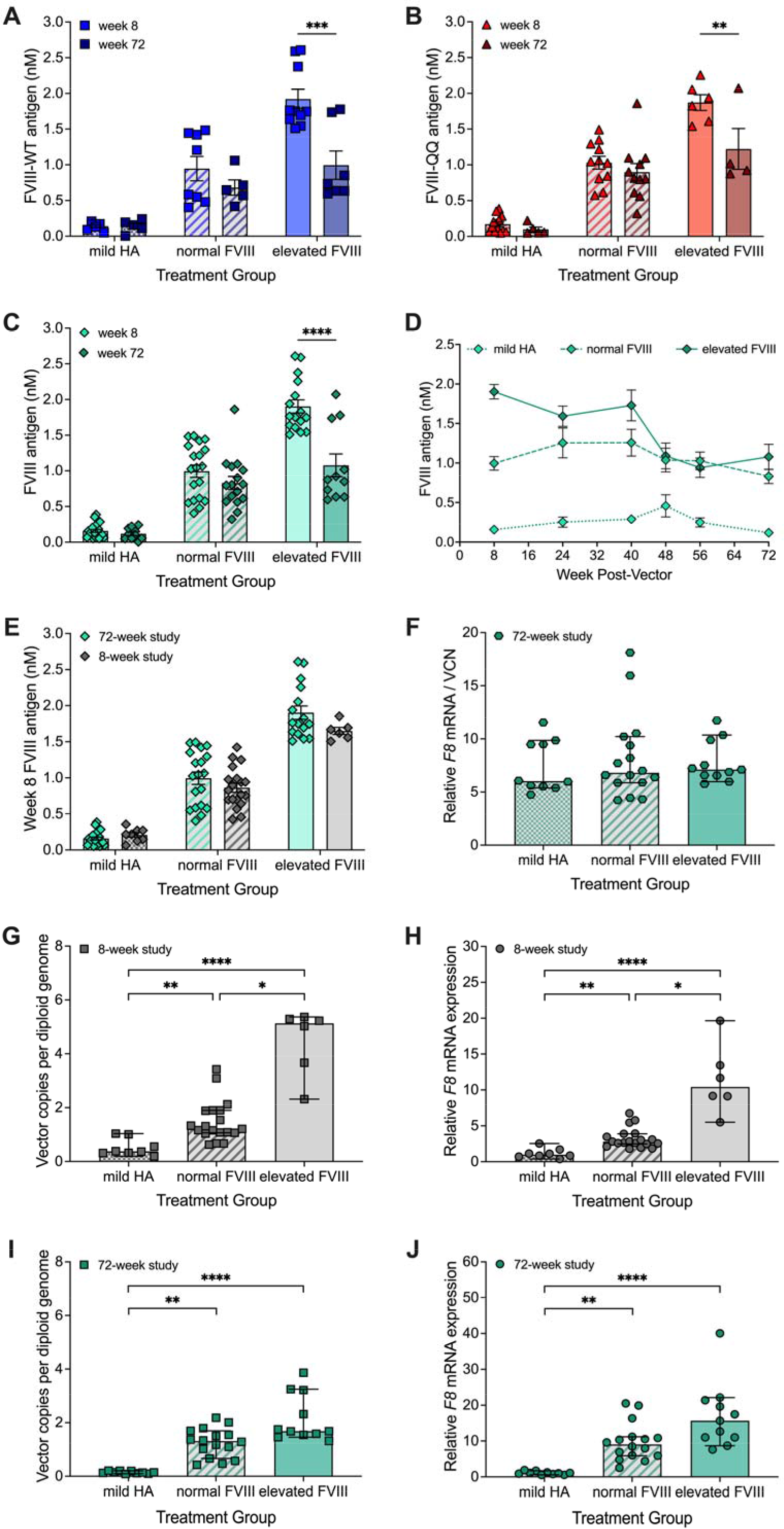
Durable FVIII expression is dependent plasma antigen levels. (**a**,**b**) HA/CD4KO mice were treated with AAV vector to express FVIII-WT (**a**) or FVIII-QQ (**b**) and assigned to the mild HA (0.05 to <0.4 nM FVIII), normal FVIII (≥0.4 to 1.5 nM FVIII), and elevated FVIII (≥1.5 to 3 nM FVIII) cohorts according to plasma antigen values measured 8 weeks post-vector. Differences in plasma FVIII levels measured at 8- and 72- weeks post-vector are shown for each antigen cohort. Significance was determined by mixed-effects analysis with Šídák’s multiple comparisons test. Individual data points with mean ± SEM are shown. (**c**) Plasma antigen at 8 versus 72 weeks post-vector for combined FVIII-WT and FVIII-QQ antigen cohorts. Differences in FVIII antigen at week 72 versus week 8 were determined by mixed-effects analysis with Šídák’s multiple comparisons test. Individual data points with mean ± SEM are shown. (**d**) Plasma FVIII antigen (mean ± SEM) over time for mild HA, normal, and elevated FVIII cohorts (n = 21, 19, and 16, respectively). (**e**) Plasma FVIII antigen (mean ± SEM) measured 8 weeks post-vector did not differ between 8-week and 72-week study mice (multiple unpaired t-tests with Welch correction for variance and Holm-Šídák correction for multiple comparisons). (**f**) Ratios of relative *F8* mRNA expression versus VCN for 72-week livers did not differ among the treatment groups (Kruskal Wallis test with Dunn’s multiple comparisons test). Individual data points with median ± 95% CI are shown. (**g**-**j**) Isolated liver VCN (**g**,**i**) and *F8* mRNA expression (**h**,**j**) for 8-week mice (**g**,**h**) and 72-week mice (**i**,**j**). VCN (**g**) and *F8* mRNA (**h**) significantly increased with cohort FVIII antigen level for 8- week mice. VCN (**i**) and *F8* mRNA (**j**) did not differ between the normal and elevated FVIII cohorts for 72-week mice. Adjusted p-values were obtained via Kruskal Wallis tests with Dunn’s multiple comparisons testing. Individual data points with median ± 95% CI are shown. *, p<0.05; **, p<0.01; ***, p<0.001; ****, p<0.0001.

Leading mechanistic hypotheses to explain declines in FVIII expression include gene silencing or vector properties that may preclude stable episome formation (*e.g.* cassette sizes and manufacturing platform^26–29^). Single AAV8 vector preparations to express FVIII-QQ or FVIII-WT were used for all expression cohorts. The oversized cassette size (5kb) demonstrated expected and comparable genomic heterogeneity between the FVIII-QQ and FVIII-WT vectors (**Extended Data Fig. 1**a,b). Significant reductions in plasma FVIII antigen were isolated to the elevated FVIII cohorts, suggesting that virion genomic heterogeneity or manufacturing platform do not mechanistically account for loss of FVIII expression. Additionally, declines in FVIII plasma antigen occurred >6 months post vector, which is beyond the window of stable episome formation demonstrated with similarly sized cassettes^27^. Overall, these data do not implicate vector properties or unstable episome formation as the cause of concentration-dependent loss of FVIII expression.

To further investigate the mechanism of FVIII expression loss, livers were harvested from mice at week 72 and analyzed for vector copy number (VCN) and *F8* mRNA. Additional HA/CD4KO mice were treated with the same AAV8 vector constructs to express FVIII-WT and FVIII-QQ in antigen matched cohorts and livers were harvested at week 8 post vector. Week 8 FVIII antigen values for each FVIII cohort did not significantly differ among mice with livers harvested at 8 versus 72 weeks post vector (**Fig. 1e**). Inconsistent with gene silencing, the ratio of *F8* mRNA to VCN did not significantly differ among the FVIII antigen groups for week 72 (**Fig. 1f**) or week 8 (data not shown) analyzed livers. As expected^27,30^, VCN and *F8* mRNA in week 8 harvested livers significantly increased by plasma FVIII antigen cohort level (**Fig. 1g,h**), which differs from small cohort observations of HA human liver biopsies post AAV vector ^31^. In contrast with week 8 data, analysis of week 72 harvested livers demonstrated no significant difference in VCN and *F8* mRNA between the normal and elevated FVIII cohorts (**Fig. 1i**,**j**), which is consistent with no significant difference in FVIII antigen between the normal and elevated FVIII cohorts at week 72 (**Fig. 1c**). These data support that decreases in FVIII expression in the elevated FVIII cohort are due to loss of VCN over time.

Given the established relationship between FVIII expression and induction of an endoplasmic reticulum (ER) stress response^30,32–34^ that can lead to apoptosis and loss of transduced cells, we investigated whether the reduction of FVIII plasma antigen in the elevated cohort was due to a cellular stress response. Markers of cellular stress, binding immunoglobulin protein (*BiP*) and C/EBP (*CHOP*) mRNA, were quantified from week 8 and 72 harvested livers. Among week 8 livers, *BiP* and *CHOP* values did not significantly differ from time-matched WT/CD4KO mice (**Extended Data Fig. 1**c), but *CHOP* expression was significantly higher in the elevated FVIII cohort relative to the mild HA and normal FVIII cohorts (**Extended Data Fig. 1**d). For week 72 analyzed livers, *BiP* was significantly elevated among all cohorts (**Extended Data Fig. 1**e) and *CHOP* was significantly elevated in the normal and elevated FVIII cohorts (**Extended Data Fig. 1**f) relative to time-matched WT/CD4KO livers. These studies established a positive association between plasma FVIII antigen and markers of cellular stress that suggest ER stress may be responsible for loss of transduced cells and, thus, VCN. Whether a competent immune system, not represented in these studies, may modulate ER stress is unclear and will be important to determine in future investigation. Overall, we conclude that FVIII expression-level dependent decreases in plasma FVIII are due to decreased VCN over time. These studies provide strong rationale to use a gain-of-function FVIII variant to restore FVIII function at low plasma FVIII antigen levels to permit durable expression.

### AAV gene addition of FVIII-QQ has enhanced function without impacting expression

We previously demonstrated that bypassing APC regulation with FVIII-QQ had improved hemostatic potency in recombinant protein studies^20^. To investigate if FVIII-QQ retained a hemostatic advantage over FVIII-WT in the setting of AAV-mediated gene transfer, HA/CD4KO mice were treated with AAV8 vectors to express FVIII-WT or FVIII-QQ. Animals were treated to achieve plasma FVIII-WT and FVIII-QQ antigen values of 0.05 nM (analogous to chromogenic-assay determined 5% of normal FVIII activity) that did not significantly differ (**Fig. 2a**). This expression-level was targeted because it is subtherapeutic for FVIII-WT in a tail clip assay^20^ and would permit sensitivity to discriminate between enhanced FVIII-QQ function and FVIII-WT. Blood loss in FVIII-QQ expressing mice normalized to that of WT mice while mice expressing FVIII-WT mice did not. Additionally, blood loss in FVIII-QQ expressing mice was significantly lower than FVIII-WT mice (one-tailed Mann-Whitney test, p=0.009). Thus, FVIII-QQ demonstrated greater hemostatic function than FVIII-WT after gene transfer (**Fig. 2b**).

**Figure 2.**
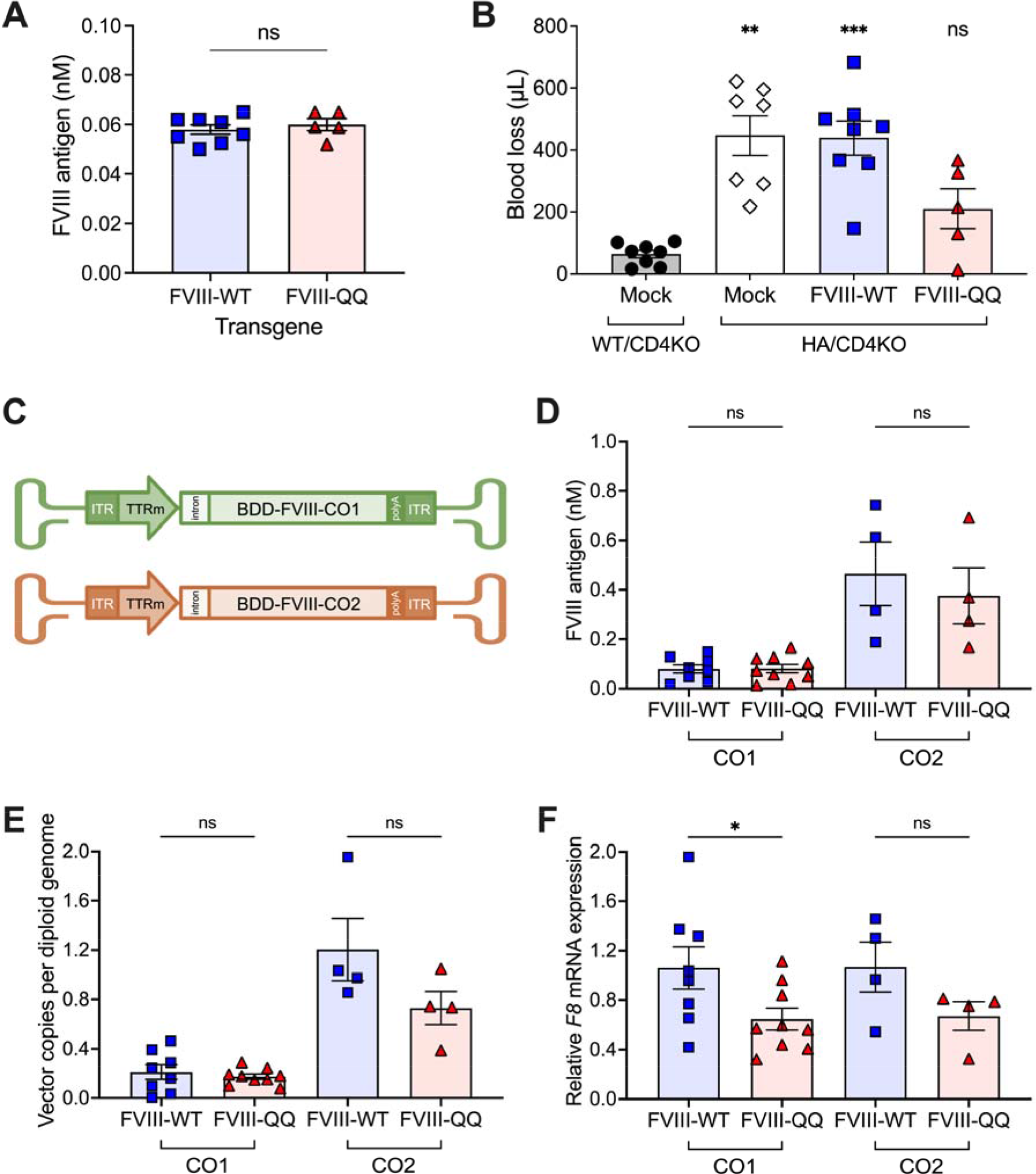
AAV gene addition of FVIII-QQ has improved *in vivo* hemostatic function relative to FVIII-WT in HA mice without impacting expression. (**a**) Week 8 plasma FVIII antigen for AAV-treated HA/CD4KO mice expressing FVIII-QQ and FVIII-WT did not significantly differ by unpaired t-test. (**b**) At 9 weeks post-vector, animals underwent tail clip assay. FVIII-QQ blood loss did not significantly differ from WT/CD4KO mice while both HA and FVIII-WT expressing mice had significantly greater blood loss than WT/CD4KO mice. Significance of blood loss versus WT/CD4KO was determined by Brown-Forsythe and Welch ANOVA tests with Dunnett’s T3 multiple comparisons test. (**c**) Adult HA/CD4KO mice were treated with AAV8 codon-optimized expression cassettes (CO1 and CO2) at 7x10^11^ vg/kg to express FVIII-QQ or FVIII-WT. Steady-state plasma FVIII antigen (**d**), isolated liver VCN (**e**), and *F8* mRNA (**f**) were analyzed for significant differences between FVIII-WT and FVIII-QQ expressing mice by unpaired t-tests. Individual data points with cohort mean ± SEM are shown. *, p<0.05; **, p<0.01; ***, p<0.001; ns, not significant.

To investigate if the introduction of the two amino acid changes of FVIII-QQ impacted expression, HA/CD4KO mice were treated with two AAV8 codon optimized constructs (CO1 or CO2) at a dose of 7x10^11^ vg/kg to express FVIII-WT or FVIII-QQ (**Fig. 2c**) in the range of mild HA (0.05 to <0.4nM), which would be targeted in clinical translation. There were no differences in week 8 steady-state FVIII-WT versus FVIII-QQ plasma concentration from either vector construct (**Fig. 2d**). Similarly, analysis of week 8-12 harvested livers demonstrated no significant difference in FVIII-WT and FVIII-QQ VCN (**Fig. 2e**) and generally no appreciable difference in *F8* mRNA (**Fig. 2f**). Lastly, consistent with no apparent differences in induction of an ER stress response, but limited by a single time point analysis, week 8 *BiP* and *CHOP* mRNA values did not differ between FVIII-QQ and FVIII-WT expressing mice (**Extended Data Fig. 2**a, b). Overall, the data do not suggest differences in FVIII-QQ or FVIII-WT expression post AAV-mediated gene transfer with codon optimized vectors.

### APC contributes to in vivo physiologic FVIIIa regulation

While acute injury models identify differences in hemostatic efficacy, they do not comprehensively recapitulate spontaneous bleeding or thrombosis^35^. To address this issue and further interrogate the role of APC in FVIIIa *in vivo* regulation, we backcrossed HA/CD4KO mice with FV-Leiden mice (FV^Q/Q^) to generate HA/FV^Q/Q^ mice that were treated with AAV vector to express FVIII-WT or FVIII-QQ (**Fig. 3a** and **Extended Data Table 1**). FV-Leiden (FV-R506Q) imparts APC resistance and is the most common genetic risk factor for venous thrombosis^36,37^. Mice homozygous for FV-Leiden exhibit a prothrombotic phenotype without diminished survival^38^. Since protein C deficiency is perinatally lethal in mice^39,40^, we hypothesized that expression of FVIII- QQ in HA/FV^Q/Q^ mice would have a major impact on the APC pathway and diminish survival.

**Figure 3:**
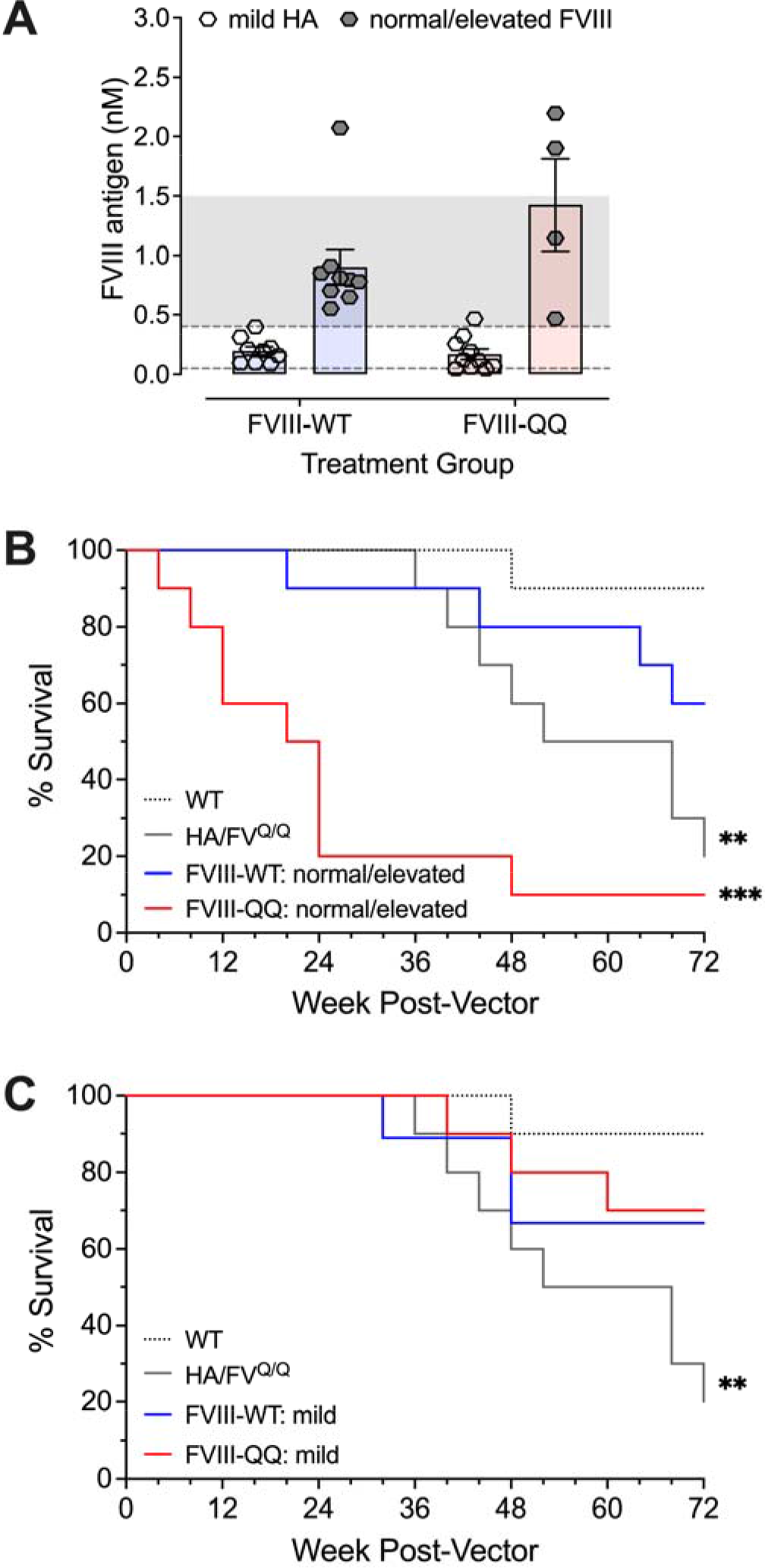
Normal to elevated plasma FVIII-QQ expression in HA/FV^Q/Q^ mice reduces survival. (**a**) HA/FV^Q/Q^/CD4KO mice were treated with AAV vector to express FVIII-WT or FVIII-QQ. Animals were assigned to mild HA (0.05 to <0.4 nM FVIII), normal FVIII (≥0.4 to 1.5 nM FVIII), and elevated FVIII (≥1.5-3 nM FVIII) cohorts according to plasma FVIII antigen 8 weeks post-vector (mean ± SEM). (**b**,**c**) Survival outcomes for mock-injected HA/FV^Q/Q^/CD4KO mice and AAV-treated HA/FV^Q/Q^/CD4KO mice expressing FVIII-WT or FVIII- QQ in the ranges of normal to elevated FVIII (**b**) and mild HA (**c**) were compared to mock-injected WT/CD4KO mice. Adjusted p-values were obtained using pairwise log-rank (Mantel-Cox) tests to determine differences in individual cohort survival versus WT/CD4KO controls and corrected for multiple comparisons using the Holm- Šídák method. **, p<0.01; ***, p<0.001.

Confirming the physiologic relevance of APC in FVIIIa regulation, expression of normal or elevated plasma FVIII-QQ antigen in HA/FV^Q/Q^ mice resulted in significantly reduced survival relative to WT/CD4KO mice (**Fig. 3b**). In contrast, similar levels of FVIII-WT expression were tolerated. Commensurate with FVIII prothrombotic risk being concentration-dependent, expression of FVIII-QQ below normal FVIII plasma concentrations (<0.4 nM) in HA/FV^Q/Q^ mice had no survival disadvantage (**Fig. 3c**). These data demonstrate clear evidence that APC regulation of FVIIIa has physiologic relevance and further support that FVIII-QQ has enhanced procoagulant function.

### FVIII-QQ is safe and does not exhibit a prothrombotic risk

To investigate FVIII-QQ prothrombotic risk in the absence of an underlying thrombophilia like FV- Leiden, a survival analysis was concurrently conducted on the same cohorts of mice analyzed for FVIII expression durability in **Fig. 1**. Specifically, survival of HA/CD4KO mice expressing FVIII-WT or FVIII-QQ in the range of mild HA, normal FVIII, or elevated FVIII concentrations were compared to WT/CD4KO mice (**Fig. 4a** and **Extended Data Table 1**) because HA/CD4KO mice do not survive to 72 weeks^41^. Among all cohorts of FVIII-WT or FVIII-QQ expressing mice, there was no survival disadvantage relative to WT/CD4KO controls (**Fig. 4b**-**d**). HA/CD4KO mice had significantly reduced survival commensurate with that previously observed for immune-competent HA/C57BL/6 mice^41^, supporting that the CD4KO background did not impact survival. To investigate further, week 12 and 32 d-dimer values of HA/CD4KO AAV treated mice and week 12 d-dimer values of HA/FVL^Q/Q^ treated mice were analyzed relative to thrombin injected positive control mice. Week 12 and 32 d-dimer values for all FVIII expression cohorts in HA/CD4KO and HA/FVL^Q/Q^ mice, except for HA/FVL^Q/Q^ mice expressing normal or elevated FVIII-QQ, significantly differed from thrombin injected positive d-dimer controls (**Extended Data 3**). These data correlate with survival study observations (**Fig. 3** and **4**) and support that FVIII-WT and FVIII-QQ expression were tolerated at all ranges of expression in HA/CD4KO mice.

**Figure 4:**
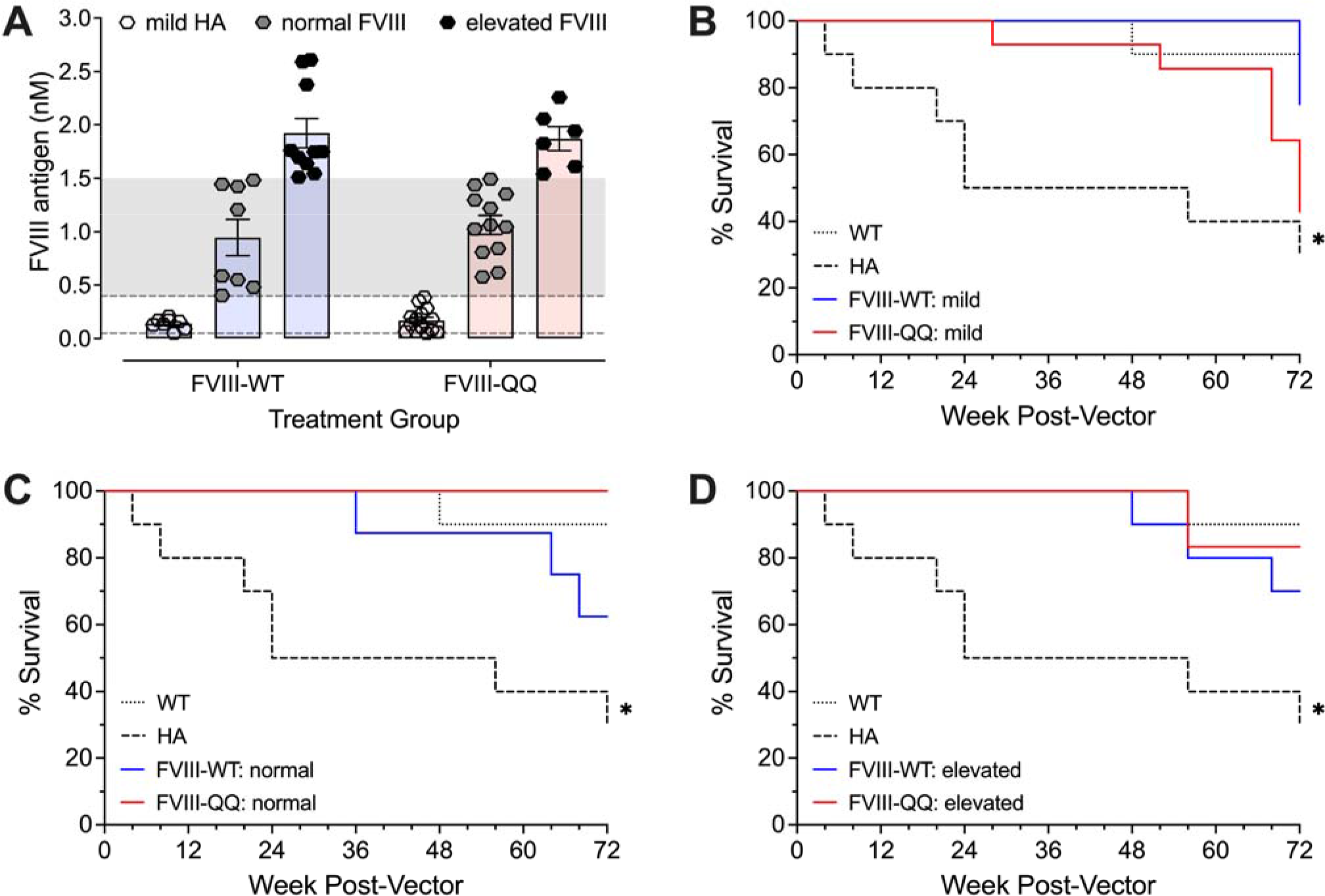
FVIII-QQ expression is tolerated in all ranges of expression in HA mice. (**a**) HA/CD4KO mice were treated with AAV vector to express FVIII-WT or FVIII-QQ. Animals were assigned to mild HA (0.05 to <0.4 nM FVIII), normal FVIII (≥0.4 to 1.5 nM FVIII), and elevated FVIII (≥1.5 to 2 nM FVIII) cohorts according to FVIII antigen (mean ± SEM) measured 8 weeks post-vector. (**b**-**d**) Survival outcomes for mock-injected HA/CD4KO mice and AAV-treated HA/CD4KO mice expressing FVIII-WT or FVIII-QQ in the ranges of mild HA (**b**), normal FVIII (**c**), and elevated FVIII (**d**) were compared to mock-injected WT/CD4KO mice. Adjusted p- values were obtained using pairwise log-rank (Mantel-Cox) tests to determine differences in individual cohort survival versus WT/CD4KO controls and corrected for multiple comparisons using the Holm-Šídák method. *, p<0.05.

To further assess prothrombotic risk *in vivo*, an Arg to Gln mutation at residue 562 was introduced by CRISPR/Cas9 to generate FVIII-QQ mice (FVIII^QQ^); wild-type mice have a Gln at position 336^42^. Homozygous female and hemizygous male FVIII^QQ^ mice were viable and fertile with normal Mendelian inheritance (**Extended Data Table 2**). Together, FVIII^QQ^ mice observations and that FVIII-QQ expression up to 2-fold normal FVIII concentrations was well-tolerated in HA/CD4KO mice demonstrates the safety of FVIII-QQ expression across two different mouse models.

### Immune responses to FVIII-QQ and FVIII-WT do not differ in immune-competent mice

To assess the immunological risk, we investigated whether FVIII-QQ may differentially break tolerance in a mouse model of relative (h)uman FVIII-WT tolerance or induce an immune response in a hFVIII naïve model. Given first-in-human HA gene therapy efforts enroll patients with established FVIII-WT tolerance^6,43^, the risk of FVIII-QQ versus FVIII-WT to break hFVIII-WT tolerance was investigated first. These studies were conducted in an immune-competent HA mouse model with ectopic hFVIII-WT expression in platelets (HA/pF8 mice)^44^ that permits relative hFVIII tolerance^45^. Mice were challenged with weekly 1µg of FVIII-WT or FVIII-QQ protein for 6 weeks followed by a larger dose of 5µg (**Fig. 5a**). FVIII-QQ and FVIII-WT challenged mice demonstrated no significant difference in the incidence or magnitude of functional FVIII inhibition determined by Bethesda titer (**Fig. 5b**) or total anti-FVIII IgG (**Fig. 5c**). Next, HA/pF8 mice without inhibitory antibodies following 4 weekly 1µg recombinant hFVIII-WT protein challenges were treated with AAV vector at a dose that conferred 1nM steady-state FVIII expression in HA/CD4KO mice. As with recombinant protein challenge, there was no significant difference in the incidence or magnitude of Bethesda inhibitory antibody titer between FVIII- QQ and FVIII-WT expressing mice (**Fig. 5d**). Commensurate with the ability of liver-directed gene transfer to induce tolerance^46,47^, all inhibitor-positive AAV-treated HA/pF8 mice tolerized (**Fig. 5d**). Consistent with prior work^25,48,49^, studies in immune-competent HA/C57BL/6 mice challenged with recombinant FVIII (**Extended Data Fig. 4**a-c) or AAV-mediated FVIII expression (**Extended Data Fig. 4**d) demonstrated a robust immune response. However, like HA/pF8 observations, there were no differences in the incidence or magnitude of inhibitor development in FVIII-QQ versus FVIII-WT treated HA/C57BL/6J mice (**Extended data Fig 4**). While limited to measuring an anti-FVIII humoral immune response only, together, these studies support that FVIII- QQ does not have increased risk of breaking FVIII tolerance or differential antibody formation relative to FVIII- WT.

**Figure 5:**
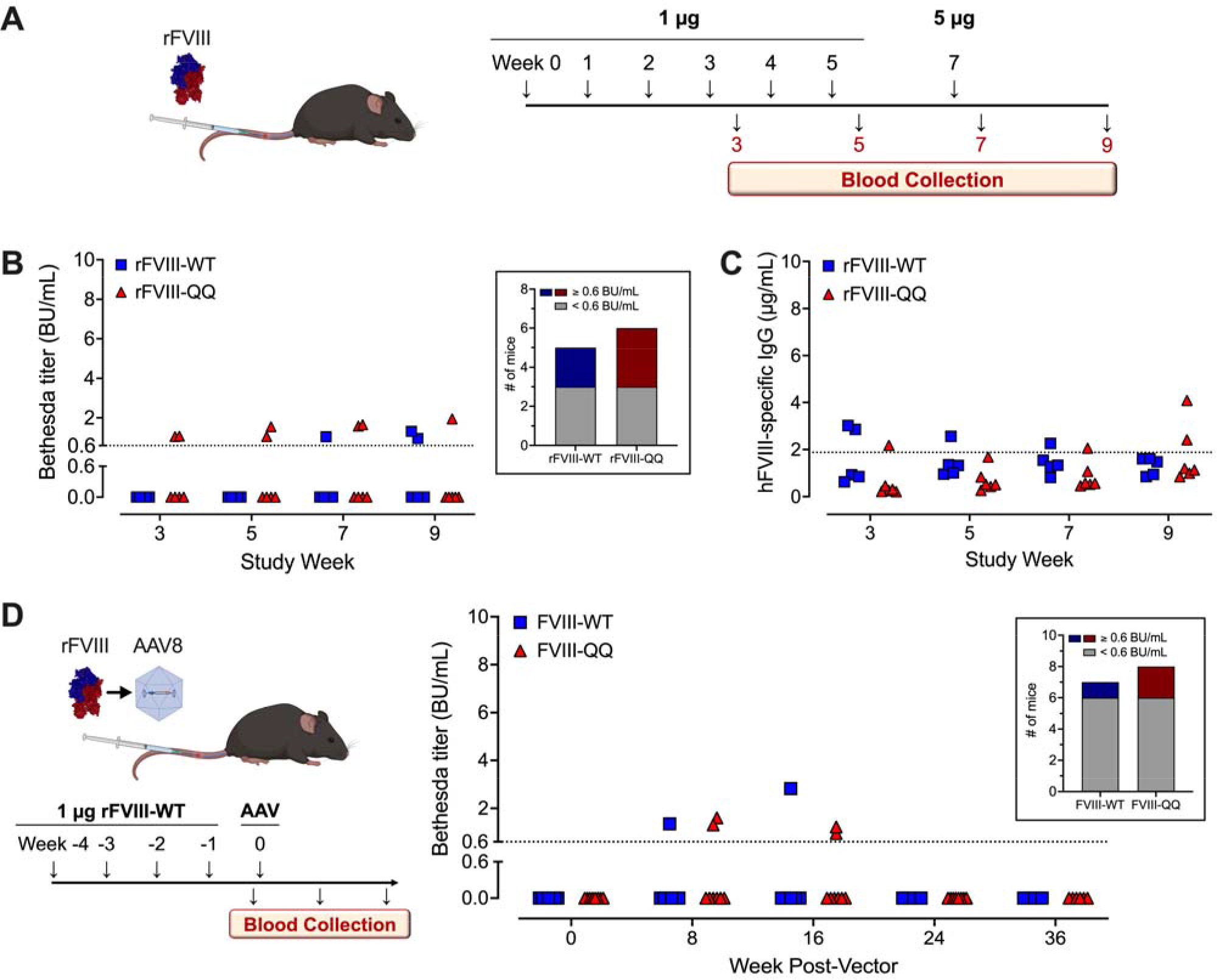
FVIII-QQ did not differentially break human FVIII tolerance relative to FVIII-WT. (**a**) Schematic of recombinant FVIII immunology study design. Bethesda (**b**) and total anti-human FVIII IgG (**c**) titers for HA/pF8 mice immunized with recombinant human FVIII-WT (n=5) or FVIII-QQ (n=6). Magnitudes of FVIII-QQ versus FVIII-WT Bethesda and IgG titers did not significantly differ (multiple Mann-Whitney tests with Holm-Šídák correction for multiple comparisons). Cumulative incidence of inhibitor development (inset in **b**) did not significantly differ between FVIII-WT and FVIII-QQ immunized mice (Fisher’s exact test). Positive Bethesda titers were defined as ≥0.6 BU/mL (**b**, dotted line). Positive IgG titers were defined as those above untreated HA/pF8 background (2 µg/mL; **c**, dotted line). (**d**) Inhibitor-negative HA/pF8 mice following recombinant FVIII- WT protein challenge were infused with AAV vector to express human FVIII-WT (n=7) or FVIII-QQ (n=8) at normal FVIII antigen levels. Inhibitor development was determined by Bethesda assay with positive titers defined as ≥0.6 BU/mL (dotted line). Bethesda titers and cumulative incidence of inhibitor development (inset) did not differ between FVIII-WT and FVIII-QQ expressing mice (multiple Mann-Whitney tests with Holm-Šídák correction for multiple comparisons and Fisher’s exact test, respectively).

### Functional assessment of FVIII-QQ

Activation of protein C requires endothelial transmembrane bound proteins (thrombomodulin and endothelial protein C receptor), which are not present in plasma^50^. Thus, no appreciable APC is generated in traditional clotting assays. As a result, the specific activities of FVIII-QQ and FVIII-WT do not differ^20^ in the absence of the PC pathway. To that end, to assess the hemostatic potential of FVIII-QQ more closely, modification of the traditional clotting assay with the incorporation of APC is required. Ultimately, this could prove beneficial or may be necessary for clinical translation of FVIII-QQ to both inform and assess clinical outcomes and safety. To investigate this, FVIII procoagulant function was assayed by activated partial thromboplastin time (aPTT) with or without APC to generate an APC resistance ratio (APCR). This approach is analogous to assays previously established in clinical practice to diagnose heterozygous FV-Leiden^51,52^. An APCR is the ratio of the clot times in the presence versus absence of APC such that a lower APCR indicates APC resistance. Plasma from AAV-treated HA (**Fig. 6a**) or HA/FV^Q/Q^ (**Fig. 6b**) survival study mice was pooled and assayed across a range of plasma FVIII concentrations. As expected, FVIII APCR values were lower for FVIII-QQ versus FVIII-WT, and APCR values for mice on the FV^Q/Q^ background (**Fig. 6b**) were lower than HA mice (**Fig. 6a**). Similarly, APCR values for a humanized system of recombinant FVIII-QQ reconstituted in human HA plasma were lower than those for equivalent concentrations of FVIII-WT (**Fig. 6c**). Consistent with prior studies of the impact of FVIII concentration on APC resistance^53,54^, APCR inversely correlated with FVIII concentration for all conditions tested. Importantly, survival outcomes for AAV-treated HA and HA/FV^Q/Q^ mice (**Fig. 3** and **Fig. 4**) correlated with APCR when stratified by FVIII concentration (**Fig. 6a**,**b**), which demonstrates that the safety of FVIII-QQ expression is APCR-dependent.

**Figure 6:**
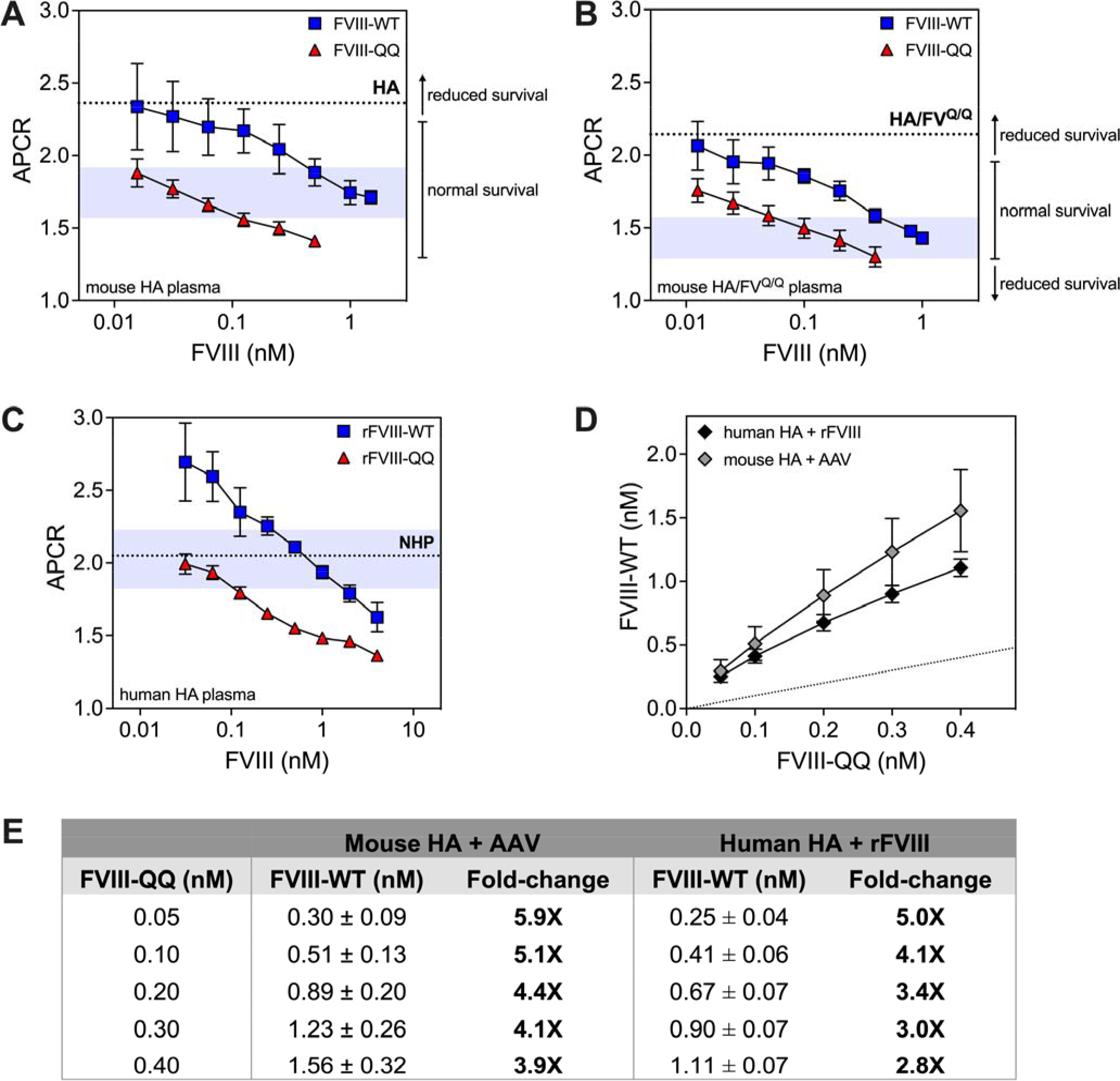
Functional assessment of FVIII *in vitro* activity in the presence of APC. (**a**,**b**) Pooled plasma from HA/CD4KO (**a**) and HA/FV^Q/Q^/CD4KO(**b**) mice expressing FVIII-WT or FVIII-QQ was assayed at varying plasma FVIII concentrations for APC resistance (APCR). The blue shaded regions represent APCR ± 10% for 1 nM FVIII-WT in the associated mouse plasma. Horizontal dotted lines represent APCR of mouse plasma without FVIII. Observed survival outcomes for varying levels of FVIII expression in HA (**a** and Fig. 4) and HA/FV^Q/Q^ (**b** and Fig. 3) mice are noted compared to their corresponding APCR values. (**c**) APCR for recombinant human FVIII-WT or FVIII-QQ reconstituted in human HA plasma at varying FVIII concentrations. The blue shaded region represents APCR ± 10% for pooled normal human plasma (NHP; dotted line). (**d**) The concentration of FVIII-WT necessary to achieve the same clot time or procoagulant activity as FVIII-QQ in the presence of APC was calculated for FVIII-QQ concentrations in the range of mild HA. The dotted line represents the line of unity. (**e**) Fold-improved procoagulant function of FVIII-QQ versus FVIII-WT in the presence of APC was calculated. Data represent mean ± SEM of 3 independent measurements performed in duplicate.

Within the translationally targeted range of FVIII-QQ expression (0.05 to ≤0.4 nM), 3-5-fold higher FVIII- WT concentrations were required to achieve the same procoagulant activity as FVIII-QQ in the presence of APC (**Fig. 6d,e**). This observation was maintained in plasma from AAV treated mice and in a fully humanized system of human HA plasma reconstituted with recombinant FVIII-WT or FVIII-QQ protein. These data are consistent with the previously demonstrated 4-5-fold improved *in vivo* hemostatic function of FVIII-QQ over FVIII-WT in multiple hemostatic injury models^20^. Thus, this aPTT-based FVIII activity assay measured in the presence of APC (“APC FVIII activity”) appears to be predictive of FVIII-QQ *in vivo* hemostatic function and it could be used as a simple clinical assay to assess FVIII-QQ function post gene transfer.

## Discussion

Despite significant progress in HA gene therapy, the goal of sustained FVIII expression that normalizes hemostasis has not been achieved. Specifically, one major limitation in the field is durability of FVIII expression, a problem that has blunted licensed HA gene therapy uptake by patients. Data in mice herein support that plasma FVIII level-dependent declines in expression are due to a reduction in VCN. These important findings address a major mechanistic gap in the field and suggest that there may be intractable barriers to expressing FVIII in hepatocytes above a certain threshold. To circumvent this, we show that FVIII- QQ is safe and has superior hemostatic function relative to FVIII-WT in the context of gene therapy. These exciting data are the first adaptation of a gain-of-function FVIII variant to demonstrate a more potent *in vivo* hemostatic response using a gene therapy approach. Our data suggest that FVIII-QQ would permit the use of lower AAV vector doses to minimize dose-dependent AAV immune toxicities, including those that limit efficacy^6,13,55^, while normalizing FVIII function at below normal plasma FVIII concentrations to permit durable expression^6,9^ and substantively advance HA gene therapy.

Unexplained year-over-year loss of initially normal FVIII-WT expression^7,11,14^ versus durable low-level FVIII-WT expression have each been demonstrated in multiple independent trials^6,9,14^. Our study is the first to demonstrate longitudinal pharmacokinetics of expression-level dependent loss of FVIII post AAV gene therapy in an animal model that are qualitatively consistent with human observations. In mice with initially elevated steady-state FVIII levels, plasma FVIII and VCN were significantly reduced 72 weeks post-vector, supporting that VCN loss is the cause of declining FVIII levels. At present, the mechanism of VCN loss is unclear, but some possibilities can be ruled out. For example, the same AAV8 vector preparations were used for all animals followed for 72 weeks, which suggests against capsid-specific epigenetic changes to expression^56^. Further, vector properties or manufacturing platform have been suggested to impact hepatocyte transgene expression and durability^29^. While this hypothesis is not supported by the data, it cannot be excluded in the absence of studies investigating dose-dependent vector toxicity. Additionally, adult mice were treated and declining FVIII plasma antigen in the elevated FVIII cohort began >40 weeks post-vector. This is outside the window of expected stable episome formation^27^ and demonstrated mouse hepatocyte proliferation^57^, suggesting against a mechanism involving either processes. Consistent with prior studies of FVIII expression in mammalian expression systems^30,32–34^, we observed a significant positive correlation between ER stress markers and FVIII expression. While these data do not provide a direct mechanistic link between ER stress and loss of VCN and interpretation is limited by possibly insensitive timing of liver sampling, we hypothesize that loss of VCN is due to FVIII expression level-dependent induction of an unfolded protein response resulting in loss of transduced cells. An ongoing study of human liver biopsy specimens post valoctocogene roxaparvovec^58^ may determine if our studies in mice translate to humans. Nonetheless, human observations expressing FVIII, including clinical trial data of an enhanced secretion FVIII variant^9^, have not achieved sustained normal FVIII expression suggesting this may not be possible with current approaches targeting hepatocyte expression. Alternatively, the goal of sustained normalized FVIII hemostatic function may best be achieved using a gain-of-function variant, such as FVIII-QQ.

To this end, we demonstrate that FVIII-QQ has greater function relative to FVIII-WT in the setting of gene therapy in multiple model systems. Interestingly, kinetic studies of FVIII/FVIIIa regulation have reported APC cleavage is up to an order-of-magnitude slower than A2-domain dissociation^59–61^, which calls into question the physiologic relevance of APC in FVIIIa regulation. Importantly, however, there are limitations of modeling FVIIIa inactivation *in vitro* due to the difficulty of purified systems to accurately replicate physiologic inter- and intramolecular interactions that are known to impact A2 dissociation^62–64^ and APC cleavage^65,66^. For the clinical translation of FVIII-QQ, this highlights the importance of *in vivo* assessment of APC on FVIIIa regulation. We demonstrated inhibiting APC regulation of FVIIIa via FVIII-QQ improves *in vivo* hemostatic function in recombinant protein studies^20^ and post gene transfer. Furthermore, induction of APC anticoagulant deficiency by expressing FVIII-QQ in HA/FVL^Q/Q^ mice recapitulated the lethal phenotype observed in protein C-deficient mice and humans^39,40,67^. This unambiguously demonstrates that APC regulation of FVIIIa has physiologic significance and extends mechanistic understanding of *in vivo* FVIIIa regulation.

The prohemostatic effect of bypassing APC regulation of FVIIIa must be balanced and thoughtfully considered with established correlations between elevated FVIII levels and venous thrombosis risk^53,68,69^. That variable FVIII expression has been observed in multiple HA gene therapy trials^8,43,70^ and excess FVIII and von Willebrand factor function are the most prothrombotic of all procoagulant proteins^68^ highlights the importance of ensuring FVIII expression is maintained within a safe therapeutic window. Indeed, the range of tolerated FVIII expression will narrow inversely with the degree of enhanced FVIII hemostatic effect, raising the possibility of diminishing therapeutic returns of efforts to improve FVIII function. Analogous to our approach, preliminary clinical trial data of a recombinant protein partially inhibiting APC anticoagulant function^71^ demonstrated marked improvement in bleeding without prothrombotic events^72^. Our studies show that all ranges of FVIII-QQ expression, including up to 2-fold normal values, were well tolerated in HA/CD4KO mice. While encouraging, we acknowledge that mice do not comprehensively recapitulate human venous thrombosis^35,73^. To address this, we also evaluated FVIII-QQ prothrombotic risk in a FV-Leiden mouse model, a provocative positive control that probes the same regulatory pathway and has a known venous thrombosis risk in humans and mice^36–38^. We found that FVIII-QQ expression in the range of mild HA (0.05 to <0.4 nM) was tolerated on the homozygous FV-Leiden background. These data provide compelling evidence that support FVIII-QQ safety in this targeted range of therapeutic expression, even in the setting of a major thrombophilia. Further, *in vitro* assessment of transgene-derived or recombinant FVIII-QQ demonstrated FVIII-QQ has 3 to 5-fold enhanced activity over FVIII-WT in the presence of APC; this is remarkably consistent with established *in vivo* enhanced potency of FVIII-QQ over FVIII-WT^20^. Thus, data support *in vitro* determination of FVIII activity in the presence of APC may accurately reflect *in vivo* hemostatic function and predict clinical outcome. Overall, data support that targeting FVIII-QQ expression in the range of mild HA may balance enhanced hemostatic function with thrombotic risk thereby permitting a range of safe therapeutic expression.

In conclusion, our data in mice demonstrate that plasma FVIII level-dependent declines in expression are due to loss of VCN and, thus, begin to address a major outstanding question in HA gene therapy. Data support that an enhanced function FVIII variant may be necessary for second-generation HA gene therapy to overcome current durability and efficacy challenges. Data reported herein support that FVIII-QQ expression below normal FVIII concentrations may safely restore hemostasis while permitting durability of expression and is the first to demonstrate that a gain-of-function FVIII variant has improved hemostatic function in the setting of gene therapy. These results address the molecular basis of a major limitation of HA AAV gene therapy and provide a rationally bioengineered solution.

## Acknowledgements

This work was supported by NHLBI K08-HL-146991 (L.A.G.), NHLBI T32-HL-007439 and T32-HL-007971 (A.R.S.), The Children’s Hospital of Philadelphia Cell and Gene Therapy Collaborative, and Asklepios BioPharmaceutical. The authors thank Drs. Rodney Camire, Ben Samelson-Jones, and Nabil Thalji for experimental design discussions and review of the manuscript.

## Author contributions

A.R.S., C.M.R., R.J.D., and X.L. performed experiments and analyzed the data. A.R.S. coordinated experiments. A.R.S. and L.A.G. designed experiments, analyzed data, and wrote the manuscript.

## Extended Data

**Extended Data Figure 1:**
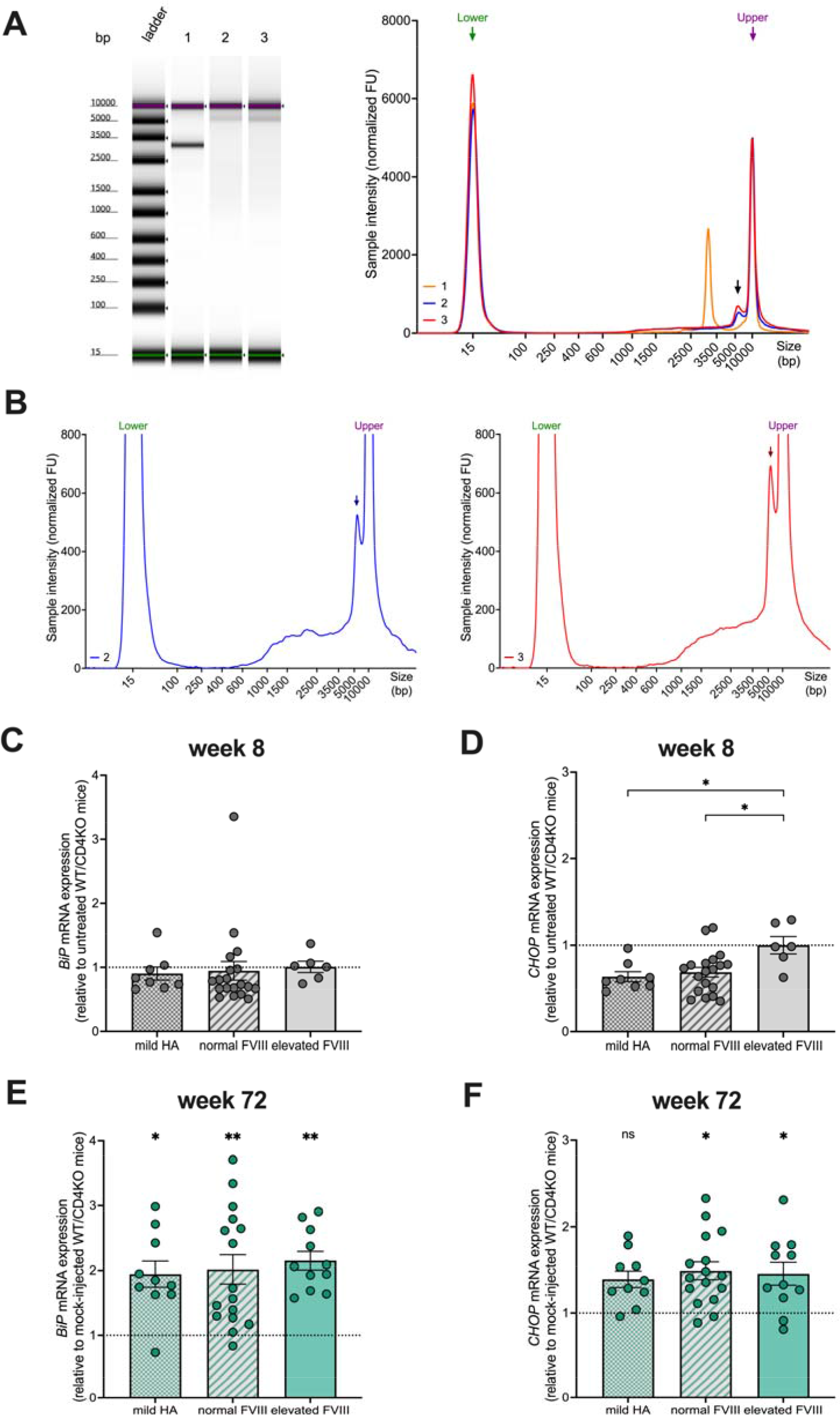
Assessment of molecular mechanisms for durable FVIII expression in HA mice. (**a**) TapeStation DNA gel of FVIII-WT (lane 2) and FVIII-QQ (lane 3) AAV8-CO1 vectors compared to control vector AAV2-CAG-eGFP (lane 1; 3 kb). Prominent 5 kb bands in lanes 2 and 3 indicate fully packaged vector genomes with smears representing heterogeneous packaging of truncated vector genomes. (**b**) Scaled signal intensities for TapeStation lanes 2 and 3. Arrows indicate fully packaged vector genomes. (**c,d**) Livers harvested 8 weeks post-vector were evaluated for *BiP* (**c**) and *CHOP* (**d**) mRNA. *BiP* expression did not significantly differ among the FVIII antigen cohorts, but *CHOP* was significantly higher for the elevated FVIII cohort compared to both mild HA and normal FVIII mice (one-way ANOVA with Tukey’s multiple comparisons test). *BiP* and *CHOP* did not differ from age-matched WT/CD4KO controls (indicated by dotted line; two-way ANOVA with Šídák’s multiple comparisons test). Individual data points with mean ± SEM are shown. (**e,f**) Livers harvested 72 weeks post-vector were evaluated for *BiP* (**e**) and *CHOP* (**f**) mRNA. Statistical significance versus age-matched WT/CD4KO controls (dotted line) are indicated (two-way ANOVA with Šídák’s multiple comparisons test). *BiP* and *CHOP* expression did not significantly differ among the FVIII antigen cohorts (one-way ANOVA with Tukey’s multiple comparisons test). Individual data points with mean ± SEM are shown. *, p<0.05; **, p<0.01; ***, p<0.001.

**Extended Data Figure 2:**
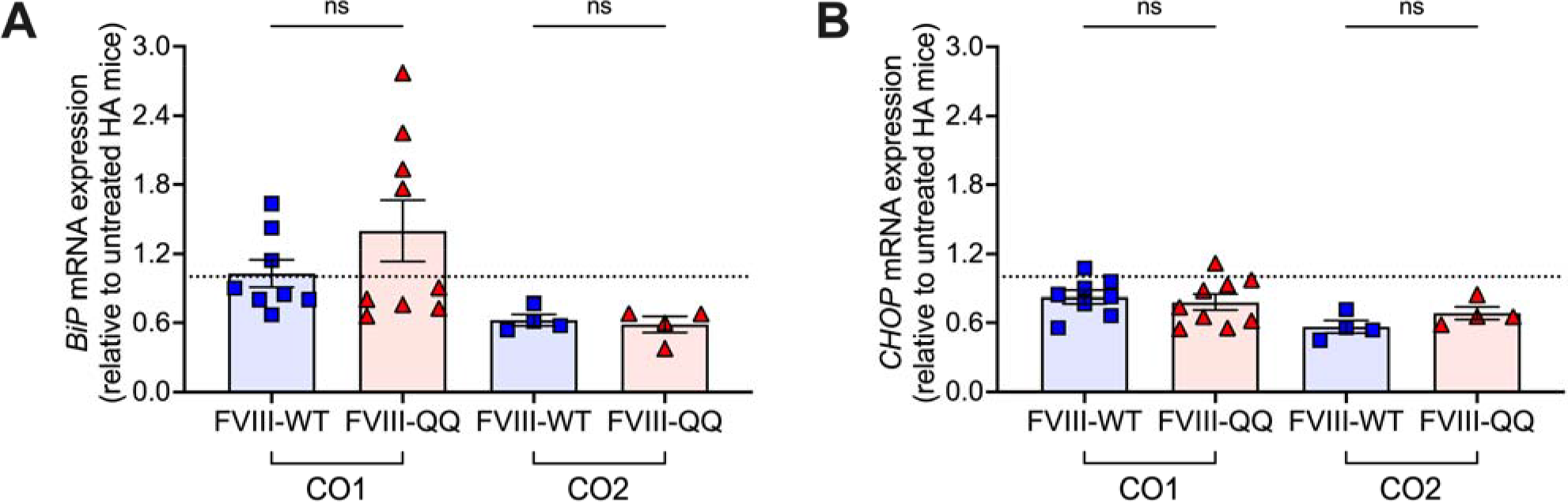
Evaluation of cellular stress in mice expressing codon-optimized *F8* transgenes. Isolated liver *BiP* (**a**) and *CHOP* (**b**) mRNA were evaluated for HA/CD4KO mice treated with 7x10^11^ vg/kg of CO1 or CO2 vector to express FVIII-WT or FVIII-QQ. There were no significant differences between FVIII-WT and FVIII-QQ mice (unpaired t-tests). Individual data points with mean ± SEM are shown. ns, not significant.

**Extended Data Figure 3:**
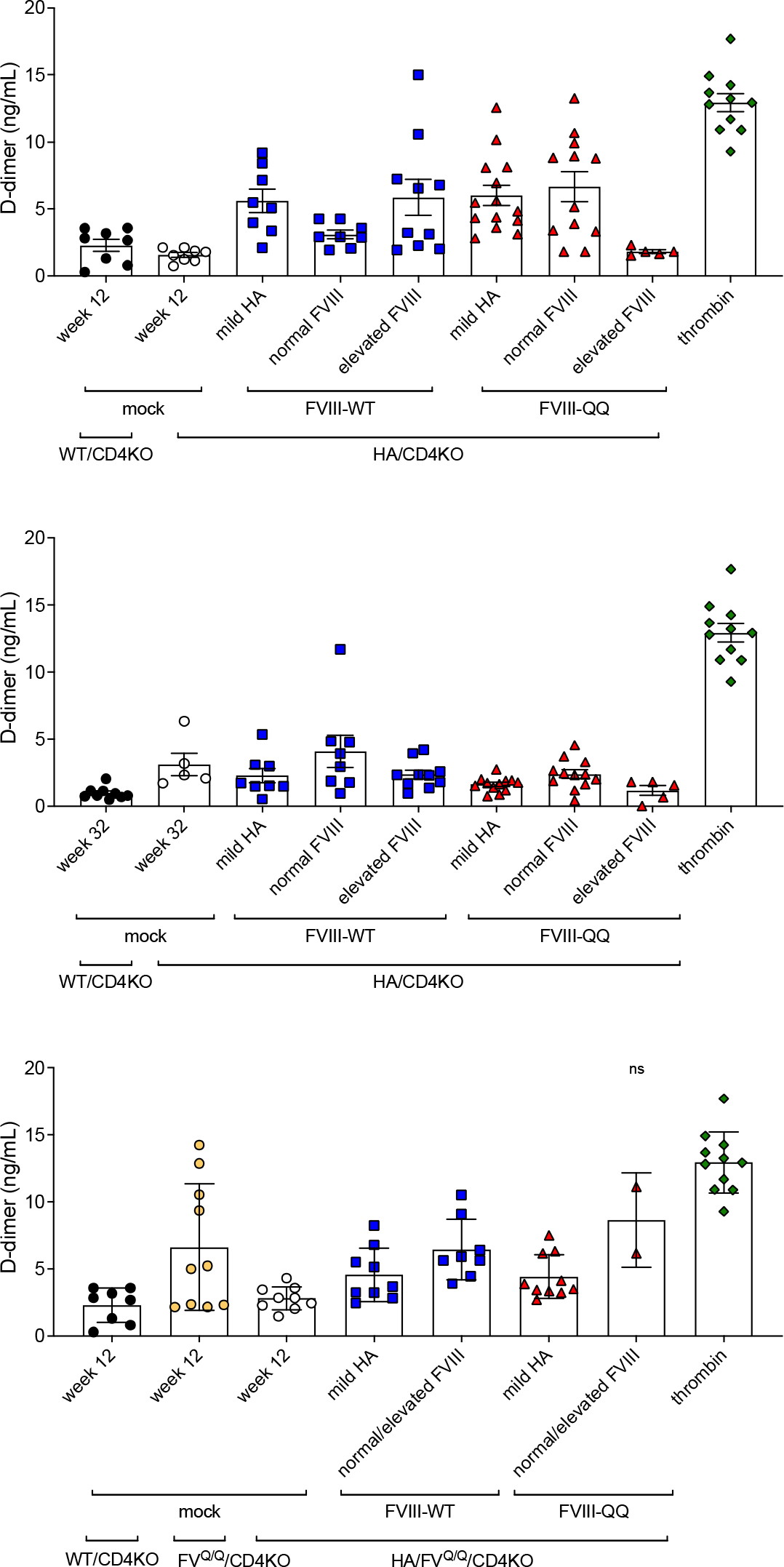
Analysis of d-dimer values of HA/CD4KO and HA/FVL^Q/Q^ mice expressing FVIII- WT and FVIII-QQ relative to positive mouse control d-dimer values. HA/CD4KO and HA/FVL^Q,Q^ mice were treated with AAV vector to express FVIII-WT or FVIII-QQ and grouped into antigen cohorts (mild HA, normal FVIII, elevated FVIII or combined normal/elevated) according to measured FVIII antigen at 8 weeks post-vector (Fig. 3a and 4a). D-dimer levels (median ± 95% CI) were measured at 12 weeks (**a**) and 32 weeks (**b**) post-vector in HA/CD4KO treated mice or at 12 weeks post vector in HA/FV^Q/Q^ treated mice (**c**). D-dimer values for all time points and FVIII expression cohorts in HA/CD4KO and HA/FVL^Q/Q^ mice, except for HA/FVL^Q/Q^ mice expressing normal or elevated FVIII-QQ, significantly differed from thrombin injected positive d-dimer controls. D-dimer values for each cohort of HA/CD4KO or HA/FV^Q/Q^ mice expressing FVIII-QQ or FVIII-WT were compared to thrombin injected mice by Brown-Forsythe and Welch ANOVA tests with test with Dunnett’s T multiple comparisons test. *, p<0.05; **, p<0.01; ***, p = 0.004, ****, p<0.0001

**Extended Data Figure 4:**
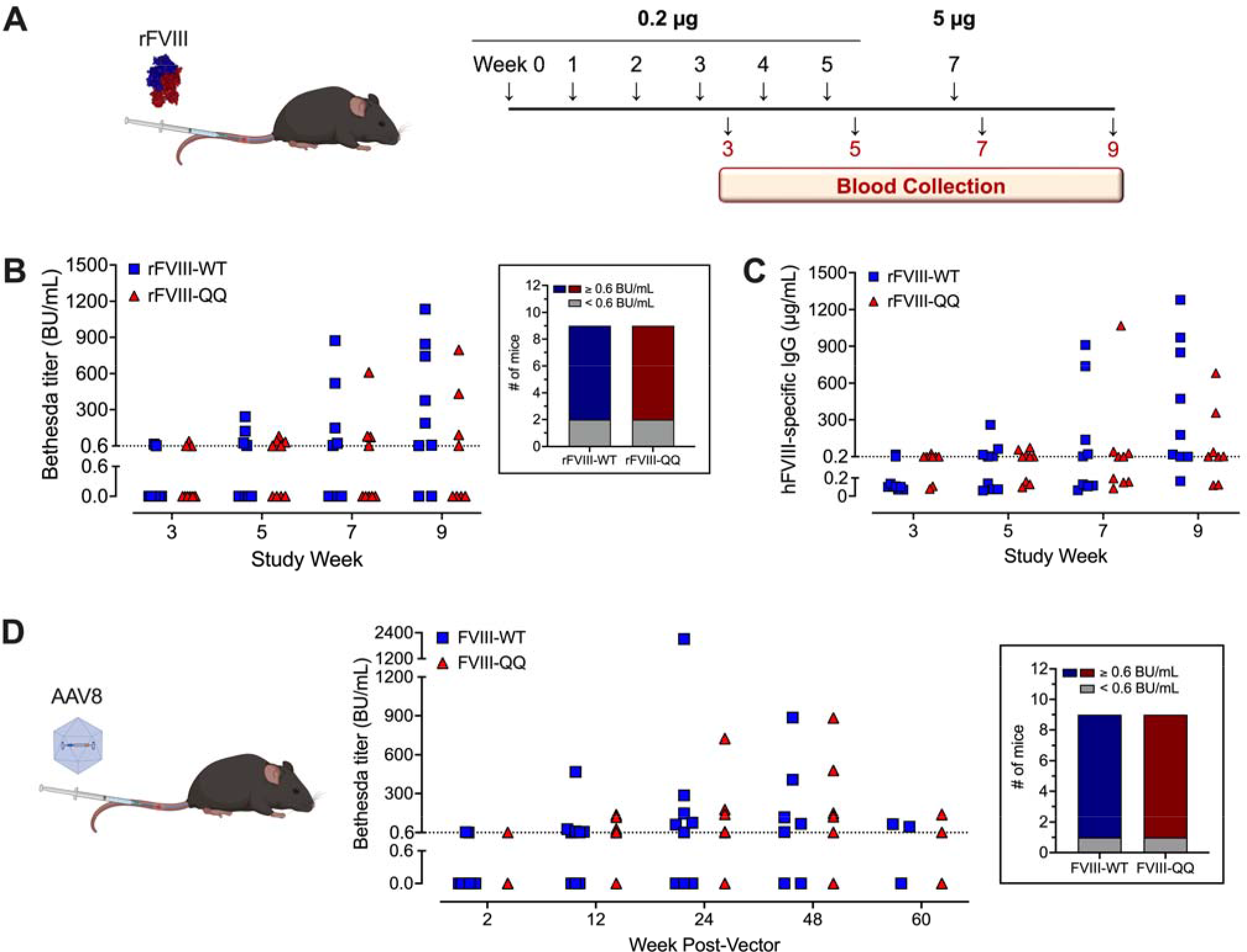
Evaluation of FVIII-WT versus FVIII-QQ immunogenicity in immune-competent HA mice. (**a**) Schematic of recombinant human FVIII immunology study design. Bethesda (**b**) and total anti-human FVIII IgG (**c**) titers for HA/C57BL/6J mice immunized with recombinant human FVIII-WT (n=9) or FVIII- QQ (n=9). Magnitudes of FVIII-QQ versus FVIII-WT Bethesda and IgG titers did not significantly differ (multiple Mann-Whitney tests with Holm-Šídák correction for multiple comparisons). Cumulative incidence of inhibitor development (inset in **b**) did not significantly differ between FVIII-WT and FVIII-QQ immunized mice (Fisher’s exact test). Positive Bethesda titers were defined as ≥0.6 BU/mL (**b**, dotted line). Positive IgG titers were defined as those above untreated HA/C57BL/6J background (0.2 µg/mL; **c**, dotted line). (**d**) HA/C57BL/6J mice were infused with AAV vector to express human FVIII-WT (n=9) or FVIII-QQ (n=9) at normal FVIII antigen levels. Inhibitor development was determined by Bethesda assay with positive titers defined as ≥0.6 BU/mL (dotted line). Bethesda titers and cumulative incidence of inhibitor development (inset) did not differ between FVIII-WT and FVIII-QQ expressing mice (multiple Mann-Whitney tests with Holm-Šídák correction for multiple comparisons and Fisher’s exact test, respectively).

**Extended Data Table 1.**
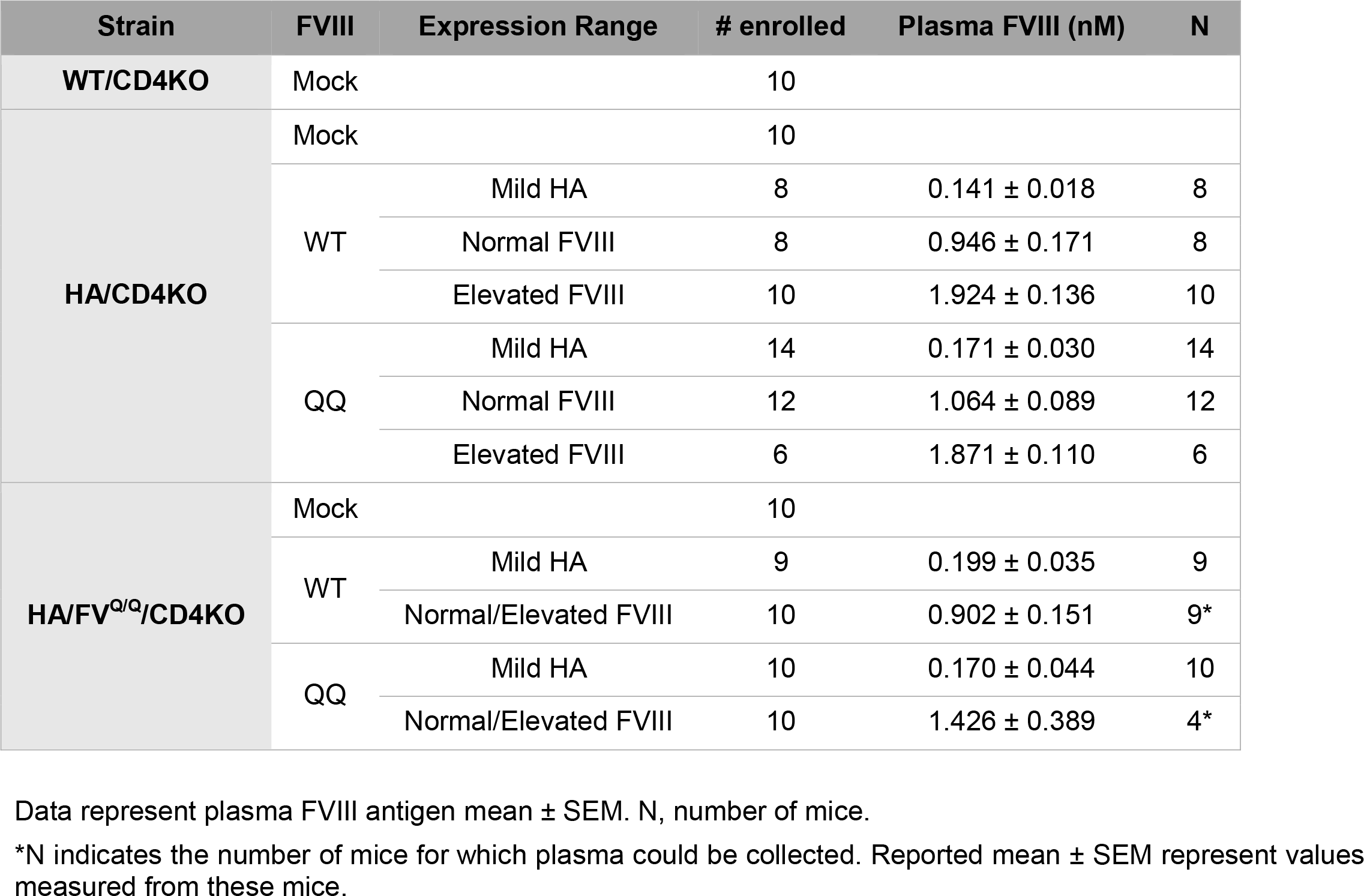
Plasma FVIII expression 8 weeks post-injection in mice followed 72 weeks for durability and survival.

**Extended Data Table 2.** Observed versus expected genetic frequencies of FVIIIQQ offspring.

|  |  | FVIII<sup>WT/WT</sup> | FVIII<sup>QQ/WT</sup> | FVIII<sup>WT/Y</sup> | FVIII<sup>QQ/Y</sup> | Total |
| --- | --- | --- | --- | --- | --- | --- |
| FVIII<sup>WT/Y</sup><br>x<br>FVIII<sup>QQ/WT</sup> | 5-9 weeks | 16 | 19 | 20 | 14 | 69 |
|  |  | 23% | 28% | 29% | 20% |  |
|  | Expected | 25% | 25% | 25% | 25% |  |
|  |  | FVIII<sup>QQ/WT</sup> | FVIII<sup>QQ/QQ</sup> | FVIII<sup>WT/Y</sup> | FVIII<sup>QQ/Y</sup> | Total |
| FVIII<sup>QQ/Y</sup><br>x<br>FVIII<sup>QQ/WT</sup> | 5-9 weeks | 12 | 19 | 22 | 18 | 71 |
|  |  | 17% | 27% | 31% | 25% |  |
|  | Expected | 25% | 25% | 25% | 25% |  |
|  |  | FVIII<sup>QQ/QQ</sup> | FVIII<sup>QQ/Y</sup> | Total |  |  |
| FVIII<sup>QQ/Y</sup><br>x<br>FVIII<sup>QQ/QQ</sup> | 5-9 weeks | 19 | 23 | 42 |  |  |
|  |  | 45% | 55% |  |  |  |
|  | Expected | 50% | 50% |  |  |  |

Chi-square and binomial testing found no significant difference between observed and expected frequencies.

